# Scaling of structural variability of ecDNA polymer condensates with copy number boosts and stabilises oncogene regulatory contacts

**DOI:** 10.64898/2026.09.25.754334

**Authors:** Mattia Conte, Sumanta Kundu, Sougata Guha, Mario Nicodemi

**Affiliations:** Dipartimento di Fisica, Università di Napoli Federico II, and INFN Napoli, Complesso Universitario di Monte Sant’Angelo, 80126 Naples, Italy

**Author notes:** Equal contribution (Mattia Conte, Sumanta Kundu).

**Keywords:** 1. ecDNA condensates, 2. oncogene regulatory contacts, 3. cell-to-cell 3D structural variability, 4. scaling behaviour, 5. polymer physics

## Abstract

Extrachromosomal DNAs (ecDNAs) form highly heterogeneous condensates in cancer cells that drive oncogene overexpression, yet how structural variability coexists with stable gene regulation remains unclear. Here, we develop a minimal polymer physics model of *MYC*-harbouring COLO320-DM ecDNAs, where BRD4-like complexes bind and bridge cognate sites along ecDNA rings. Above a critical binder concentration, ecDNAs phase separate into condensates exhibiting diverse conformations because of their thermodynamic folding degeneracy. Despite this variability, condensates retain conserved interaction scaffolds that give rise to reproducible contact patterns, including *in-trans* associated domains (I-TADs), genomic regions enriched in intermolecular regulatory contacts between distinct ecDNAs. We find that condensate 3D architecture follows universal scaling relations with ecDNA copy number, *n*, remaining robust to model parameter changes. Regulatory contacts within I-TADs increase linearly with *n*, yet they are one order of magnitude stronger than in size-matched control regions outside I-TADs, whereas their relative fluctuations are markedly suppressed as *n* increases. This scaling produces enhanced, low-noise regulatory environments for oncogenes embedded within I-TADs, such as *PVT1-MYC* fusions, whereas the canonical *MYC* copy, located outside, is less amplified as experimentally observed. Our findings reveal universal polymer physics principles underlying ecDNA condensate organization, offering a mechanistic basis for selective oncogene amplification and potential advantages in cancer progression.

**One line summary:** Despite broad single-cell 3D structural heterogeneity of ecDNA condensates, increasing ecDNA copy number enforces universal scaling laws that boost oncogene regulatory contacts within *in-trans* associated contact domains (I-TADs) and suppress their relative fluctuations, thereby stabilizing specific oncogene activation and potentially conferring a selective advantage to cancer cells.

## INTRODUCTION

Extrachromosomal DNAs (ecDNAs) are common in aggressive tumours and are associated with poor clinical outcomes^1–6^. These circular, megabase-scale DNA elements lack centromeres and telomeres, and can harbour oncogenes^3,7^ as well as regulatory elements^8,9^ that are recurrently amplified, likely conferring a selective fitness advantage^1,10^. EcDNAs are highly mobile within the cell nucleus and can occur in hundreds of copies, with substantial heterogeneities from cell to cell^11–13^. Their spatial organisation can reshape oncogene activation through contacts with other ecDNAs, chromosomal loci, and nuclear condensates, in a new paradigm of gene regulation^1,7,9,14–18^. In several cancers, such as COLO320-DM colorectal cancer cells, ecDNAs cluster into micron-sized hubs composed of dozens of rings, enhancing intermolecular enhancer-gene interactions and driving strong oncogene overexpression^8,14–16,18,19^.

EcDNAs are also a major source of intratumoral heterogeneity, a feature closely linked to uneven clinical outcomes. Distinct cells within the same tumour can differ in ecDNA content, sequence composition, chromatin state and copy number, thereby generating substantial genetic, epigenetic and structural variability^8,20–26^. Even when a single ecDNA species predominates, copy-number heterogeneity across tumour cells can arise for example by random ecDNA inheritance during cell division, coupled to adaptation to metabolic stress or treatment^1,10,27,28^. These variations can affect gene expression beyond simple dosage effects, particularly in tumours where ecDNAs form transcriptionally active hubs^15,16,18^. However, how cell-to-cell differences in ecDNA copy number influence the spatial organisation of ecDNA condensates, and how this in turn affects oncogene regulation, remains very difficult to explore at the single-molecule level with current technologies.

To address this problem, here we employ polymer physics models of ecDNAs in COLO320-DM colorectal cancer cells, whose molecular elements have been comparatively well characterised experimentally^15,16^. Most COLO320-DM ecDNAs share a common genomic architecture that includes a canonical copy of the *MYC* oncogene, two copies of a *PVT1-MYC* fusion, along with a set of cis-regulatory elements^15,16^. This makes COLO320-DM a particularly suitable system to isolate, to a first approximation, the contribution of ecDNA copy number to the 3D conformational variability of ecDNA condensates. In these cells, ecDNAs form hubs interacting with BRD4, a bromodomain and extraterminal domain (BET) family protein, and with MED1, a key transcription coactivator, driving strong overexpression of the *PVT1-MYC* fusions, but not of the canonical *MYC*^15,16^.

We use a minimal physics model in which *n* ecDNAs are represented as ring polymers confined within a defined volume, representing the nuclear space, and interacting with BRD4-like complexes that bind cognate sites along the ecDNA sequence^18^. The model was shown to undergo a phase transition as a function of binder concentration: below a critical threshold, ecDNAs are dispersed in the assigned volume, whereas above the threshold they form hubs by phase separation of polymer condensates^18^. The condensates lead to the assembly of specific *in-trans* associated contact domains (I-TADs) among ecDNAs, which constitute local environments enriched, beyond copy number effects, for regulatory contacts between oncogenes and enhancers. The genomic positions of these domains are determined by the arrangement and strength of BRD4 binding sites along the ecDNA sequence, so that only selected loci participate in the resulting *in-trans* interaction scaffold. The *PVT1-MYC* fusions, which are strongly bound by BRD4, fall within multiple I-TADs and acquire markedly enhanced regulatory interactions beyond copy number effects, whereas the canonical *MYC* copy, located outside these domains, does not, consistent with transcription data^15,18^. Notably, the ensemble-averaged contact matrix predicted by the model in the condensate state was found to match independent Hi-ChIP^15^ and ChIA-PET^16^ measurements, supporting that the model captures the essential biophysical mechanisms underlying ecDNA hub formation and structure^18^.

However, several important questions remain: how does ecDNA copy number influence the 3D conformational variability of individual ecDNAs within condensates? How can heterogeneous single-molecule folding give rise to reproducible contact patterns within and across ecDNA rings? And how do these emergent structural features affect oncogene regulatory interactions?

Using the polymer model, here we investigate the variability of the spatial organization of ecDNA condensates at the single-molecule level, and how that affects I-TADs and oncogene regulatory contacts. We find that ecDNA condensates exhibit broad 3D conformational heterogeneity, reflecting the thermodynamic degeneracy of the folded states accessible to individual ecDNA molecules and to whole condensates. As a result, single-molecule ecDNA contact and distance patterns vary substantially both *in-cis* and *in-trans*, i.e., within individual rings and between distinct rings. However, this variability does not erase the underlying organisation of the condensate. As the number of rings *n* in the condensate increases above half a dozen, the average Pearson correlation between the distance maps of two distinct ecDNAs plateaus at around r=0.5 *in-cis* and r=0.3 *in-trans*. These non-zero plateaus indicate that, although thermal fluctuations generate diverse conformations and variable contacts between distinct ecDNAs, the condensates retain a conserved and reproducible structural scaffold, characterized by statistically stable regulatory interactions.

We further show that key condensate structural features, including their volume, contact patterns, and I-TAD architecture, obey robust power-law scaling behaviours with ecDNA copy number, *n*, which do not depend on the finer details of the model, such as binder-ecDNA affinities or interaction potentials , consistent with polymer theory^29^. Contacts within I-TADs increase linearly with *n*, reflecting copy-number effects and steric constraints that limit the number of rings each ecDNA can contact. Crucially, however, I-TAD contacts are approximately an order of magnitude stronger than those in equivalently sized regions outside I-TADs. Conversely, the relative fluctuations of I-TAD contacts decrease as an inverse power of *n*, and they are markedly lower than fluctuations of contacts outside I-TADs. Thus, the regulatory contacts of genes within I-TADs are not only enriched in large condensates, but also stabilised, producing an enhanced, low-noise regulatory environment.

Finally, we assess how this structural heterogeneity and its scaling properties affect the regulatory landscape of the distinct *MYC* copies on COLO320-DM ecDNAs. We focus on a regulatory element positioned at comparable genomic distance from the canonical *MYC* copy (*MYC-1*) and from the *PVT1-MYC* fusion (*MYC-2*), which lie respectively outside and inside I-TADs. Despite their similar genomic separation, the regulatory element is consistently closer in 3D space to *MYC-2* than to *MYC-1*, both *in-cis*, when the genes lie on the same ecDNA ring (260 nm vs 350 nm), and *in-trans*, when they lie on distinct rings within the condensate (310 nm vs 440 nm for the closest neighbouring ring). That demonstrates that I-TAD membership, rather than linear genomic separation along the ecDNA sequence, is a key determinant of regulatory proximity within ecDNA condensates.

Together, these results show that high-copy-number ecDNA condensates enhance and stabilise oncogene-regulatory interactions through the conserved structural scaffolds of I-TADs and their universal scaling behaviours, despite the high conformational variability generated by thermal fluctuations. The resulting high-signal, low-noise regulatory environments may render oncogene transcription strong and persistent, offering a mechanistic basis for the observed selective evolutionary advantage of cancer cells harbouring large ecDNA copy numbers. More broadly, the scaling laws we uncover establish a quantitative link between ecDNA copy number and 3D regulatory architecture, providing a predictive framework for interpreting ecDNA-driven phenotypes across tumour contexts.

## RESULTS

### Polymer model of ecDNAs in COLO320-DM colorectal cancer cells

*MYC*-harbouring ecDNAs in COLO320-DM colorectal cancer cells are circular DNA molecules approximately 4.3 Mb in length^15,16^. They are composed of multiple fragments derived mostly from chromosomes 8, together with smaller segments from chromosomes 6, 13 and 16 (**Supplementary Fig. 1**). The chromosome 8 fragments host the canonical copy of *PVT1* and *MYC* genes, along with a broadly distributed set of BRD4 binding sites (**Fig. 1a**). Each ecDNA also carries two copies of a *PVT1-MYC* fusion, an additional *PVT1* fragment, and a collection of cis-regulatory elements. In interphase nuclei, COLO320-DM ecDNAs assemble into micron-sized hubs, each containing on average on the order of ten rings, held together by BRD4^15^ and interacting with MED1^16^, a key transcription coactivator.

**Figure 1.**
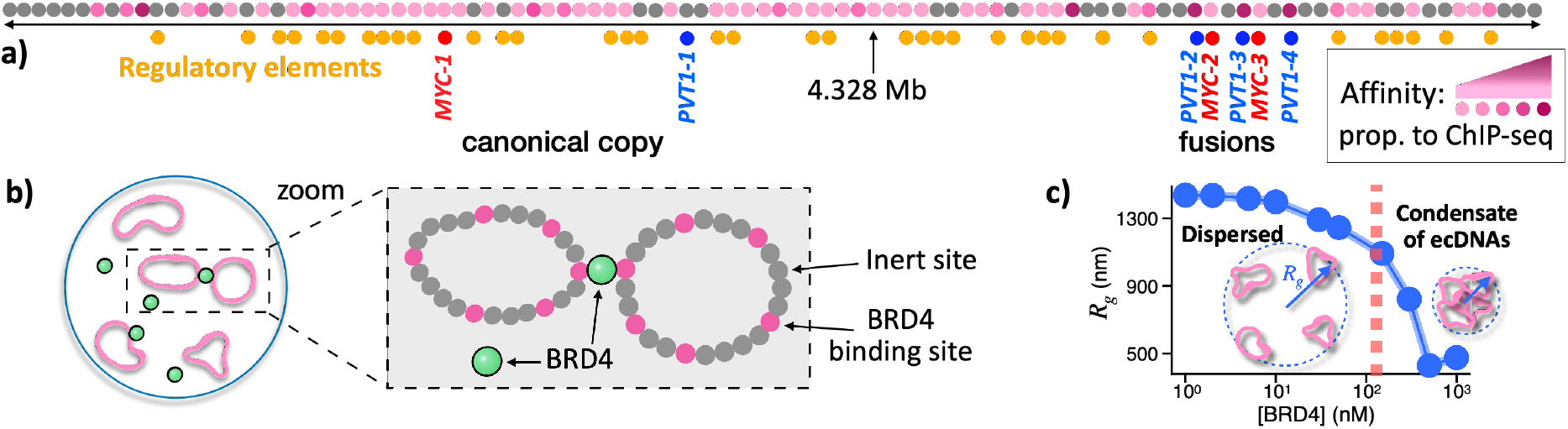
A minimal polymer model of ecDNA condensate formation in COLO320-DM. **a)** Linear scheme of the COLO320-DM ecDNA ring model, coarse-grained at 50 kb resolution. BRD4-binding sites are shown in magenta, with shade proportional to the experimental ChIP-seq signal^15^ and hence to the assigned model binding affinity. The ecDNA copies of *MYC* (*MYC-1*, *-2* and *-3*, red), *PVT1* (*PVT1-1*, *-2*, *-3*, and *-4*, blue) and cis-regulatory elements (yellow) are also mapped^15^. **b)** Schematic of the Strings and Binders (SBS) polymer model of ecDNAs in COLO320-DM. Each ecDNA is represented as a self-avoiding ring polymer confined in a given volume and composed of inert beads (grey) and BRD4-binding beads (magenta), which can be bridged by diffusing BRD4-like complexes (green). The genomic arrangement of the ecDNA chromosomal fragments and of its BRD4 binding sites is mapped at 50 kb resolution^15^. **c)** Phase transition of a representative 10-ring system as a function of BRD4 concentration. As [BRD4] increases above a threshold, ecDNA rings phase separate from a dispersed to a condensed state, as indicated by a sharp decrease in the gyration radius of the system, *R*_g_, corresponding to the formation of a shared ecDNA condensate.

To investigate the 3D structure of ecDNA condensates at the single-molecule level, we consider a polymer physics model incorporating only the essential molecular components mentioned before^18^: polymer rings bearing BRD4 binding sites and their cognate BRD4 molecular complexes. Specifically, we employ the Strings and Binders (SBS), a well-established polymer model of chromatin^30–34^. In this model, each ecDNA is represented as a self-avoiding ring polymer chain of beads, each corresponding to a 50 kb DNA window (**Fig. 1b**). BRD4 binding sites along the chain (magenta in **Fig. 1a,b**) are assigned from ChIP-seq data^15^ and represented as beads that can be bound and bridged by diffusing BRD4 molecules (green in **Fig. 1b**). Binding affinities are set to match the intensity of the corresponding ChIP-seq signals in the weak biochemical energy range (2.25-6.25 k_B_T; **Methods**), consistent with prior experimental measurements of BRD4 binding and with earlier chromatin modelling approaches^33,35–38^.

BRD4 is also known to participate in multiprotein complexes, for example through its extra-terminal (ET) domain engaging cofactors such as MED1, which can interact with one another and thereby enhance molecular stability and function^39,40^. Accordingly, in our simulations we model these complexes (green circles in **Fig. 1b**) rather than isolated BRD4 molecules, and allow them to interact with one another through a weak mutual affinity (≈3.1 k_B_T). To test the robustness of our conclusions, we further considered a range of model variants as reported in^18^ (**Methods**).

The system evolves under Brownian dynamics, which we explore through extensive Molecular Dynamics (MD) simulations. At the start of each simulation, rings and BRD4 complexes are randomly dispersed within a fixed simulation box, used as a proxy for the accessible nuclear volume, and converge to equilibrium. The number of rings, *n*, is fixed in each MD simulation. For each parameter set, we sampled up to 10^3^ independent equilibrated conformations (**Methods**). Although much of the analysis below focuses on the representative case *n*=10, consistent with the reported average size of COLO320-DM hubs^15^, we explored ecDNA copy numbers spanning one order of magnitude, from *n*=1 to *n*=20. Within each ring, we also mapped the experimentally reported copies of *MYC* (*MYC-1, -2, -3*; shown in red, **Fig. 1a**) and *PVT1* (*PVT1*-*1, -2, -3, -4*; blue), together with their cis-regulatory elements (yellow), allowing us to track how specific genes and regulatory regions interact within and across ecDNAs.

We previously showed^18^ that, when BRD4 concentration surpasses a critical threshold, ecDNA rings undergo a phase transition from a dispersed state, in which they are randomly distributed in the available volume, to a condensed state, in which they phase-separate into a single hub (**Fig. 1c**). Notably, the model transition threshold, in the range of fractions of micro-mole/l, lies close to concentrations reported in vitro for phase separation of transcriptional condensates^35^. As dictated by polymer physics^29^, these two regimes define two distinct folding classes corresponding to the system thermodynamic phases: in the dispersed state, entropic forces dominate and ecDNAs explore the available volume largely independently, whereas in the condensed state multivalent BRD4-mediated bridges between ecDNA cognate binding sites become thermodynamically favourable, lowering the free energy of clustered configurations and triggering collapse into a single condensate. This transition is marked by a sharp drop in the order parameters of the system, such as its gyration radius, *R*_g_, or the distance between rings’ mass centres, as a function of BRD4 concentration (**Fig. 1c**, **Supplementary Fig. 2**, and **Methods**). The resulting condensed state mirrors the formation of ecDNA hubs in COLO320-DM observed in recent microscopy experiments^15,16^.

Importantly, the ensemble-averaged contact maps produced by the model in the condensed regime were previously shown to recapitulate independent Hi-ChIP and ChIA-PET measurements in COLO320-DM^18^, supporting the notion that phase separation and multivalent bridging are core physical drivers of ecDNA hub formation. Our analysis therefore focuses on the non-random 3D architecture of the condensed state, which underlies the regulatory behaviour of ecDNA hubs.

### EcDNA condensates exhibit broad single-molecule structural heterogeneity yet retain a conserved contact scaffold

To quantify the 3D structural variability of ecDNA condensates at the single-molecule level, we computed *in-cis* and *in-trans* distance and contact matrices across independent equilibrium conformations of the model, corresponding to distinct realizations of single ecDNA condensates (**Fig. 2a, Methods**). Here, *in-cis* denotes contacts or distances within the same ring, whereas *in-trans* refers to those between distinct rings. We then examined how these single-molecule matrices, and their ensemble averages, vary with ecDNA copy number, *n*, across the sampled BRD4 concentrations and binding affinities.

**Figure 2.**
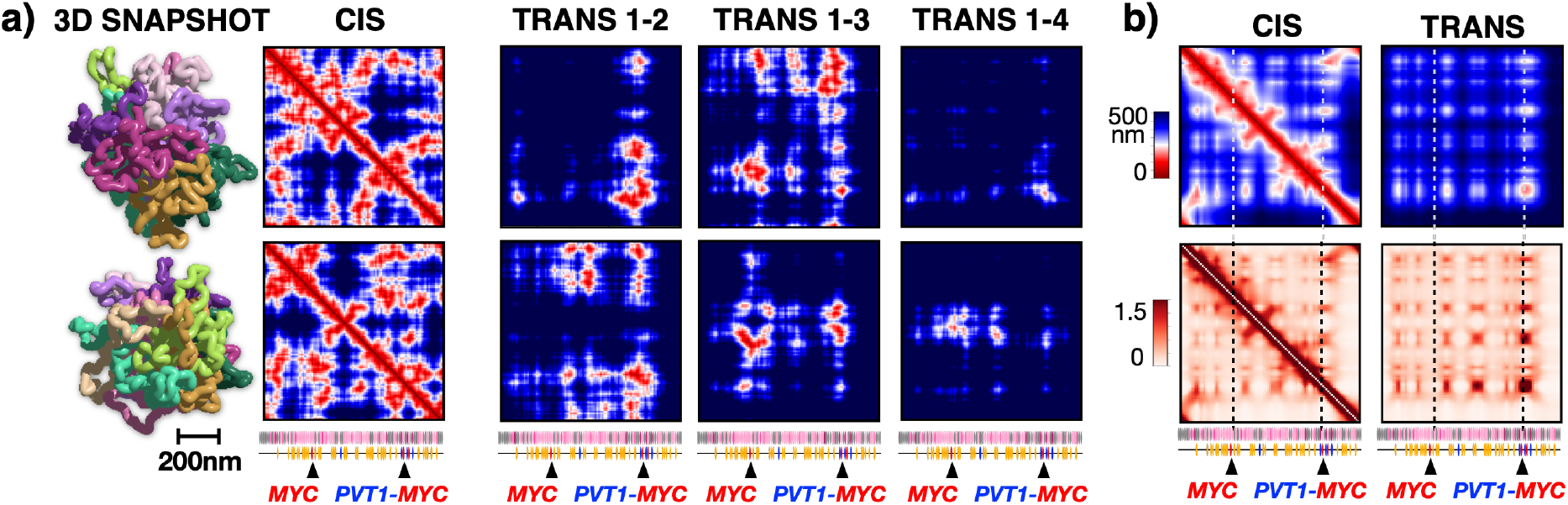
Condensates of ecDNA model rings have a broad structural heterogeneity but retain reproducible average contact patterns. **a)** Single-molecule 3D structures of ecDNA condensates, shown here for 10-ring systems, are highly heterogeneous, as illustrated by two representative conformations and by the corresponding *in-cis* and *in-trans* distance matrices of their rings. *In-cis* matrices report pairwise distances within the same ring, whereas *in-trans* matrices report those between a ring and distinct neighbouring rings. Condensate architecture breathes under thermal fluctuations, spanning a variety of distinct conformations belonging to a single conformational class. **b)** The average distance (top) and contact (bottom) matrices *in-cis* and *in-trans* per polymer are shown for a 10-ring condensate. Despite the strong single-molecule heterogeneity shown in **a)**, contacts are not random as patterns *in-cis* as well as specific *in-trans* associated contact domains (I-TADs) are formed between strong BRD4 binding sites. The *PVT1-MYC* fusion is enriched of regulatory contacts as it is part of I-TADs, whereas the canonical *MYC* is not.

In the dispersed phase, the rings randomly float as independent self-avoiding polymers in the available volume, exploring random conformations by thermal fluctuations. Accordingly, their average contact and distance maps are featureless, reflecting only transient and fleeting encounters (**Supplementary Fig. 3**). By contrast, above the phase separation threshold, multivalent BRD4-mediated bridges drive the formation of a single polymer condensate. In this regime, individual condensates display a high degree of folding with substantial conformational degeneracy, manifested in broadly heterogeneous single-ring contact and distance patterns *in-cis* and *in-trans* across distinct realisations (i.e., in single cells, **Fig. 2a**). However, specific interactions are preferentially established between regions enriched in strong BRD4 binding sites, generating reproducible patterns in the ensemble-averaged distance and contact matrices (**Fig. 2b**). These include well-defined domains in cis (TADs) and, importantly, in trans, where *in-trans* associated contact domains (I-TADs) emerge as genomic regions enriched in intermolecular contacts between distinct ecDNAs (**Supplementary Fig. 4a; Methods**).

Thus, although individual ecDNA condensates sample a broad ensemble of thermodynamically allowed conformations, condensation selects a conserved interaction scaffold, from which I-TADs and their regulatory consequences emerge.

### Scaling behaviours of ecDNA condensates

To quantify the structural variability of ecDNA condensates, we first analysed how their volume, 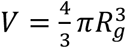, where *R_g_* is the condensate gyration radius, fluctuates across independent equilibrium configurations (**Fig. 3a**, left panel). For a fixed copy number *n*, condensate volume has a broad distribution (**Fig. 3a**, centre-left), consistent with the marked structural heterogeneity described above. Despite this variability, its average behaviour follows simple scaling laws with ecDNA copy number. The mean condensate volume, ⟨*V*⟩, increases linearly with *n*, as expected for the growth of a compact polymer assembly upon addition of further rings (**Fig. 3a**, centre-right). By contrast, the relative volume fluctuation, Δ*V*/⟨*V*⟩ = (⟨*V*^2^⟩ − ⟨*V*⟩^2^)^1/2^/⟨*V*⟩, decays to zero as *n*^-1/2^ (**Fig. 3a**, right), consistent with Central Limit expectations, whereby fluctuations arising from individual rings are progressively averaged out in larger clusters. That is in line with the general physics of topologically constrained ring-polymer assemblies, whose compact organisation and fluctuations have been extensively characterised in simulations and theory^41–48^. Overall, these scaling behaviours reveal the stabilising effect of increasing copy number and show that, once formed, condensates behave as random, yet mechanically coherent polymer aggregates whose macroscopic properties are dictated by general principles of polymer physics, rather than by the microscopic details of the model^29^.

**Figure 3.**
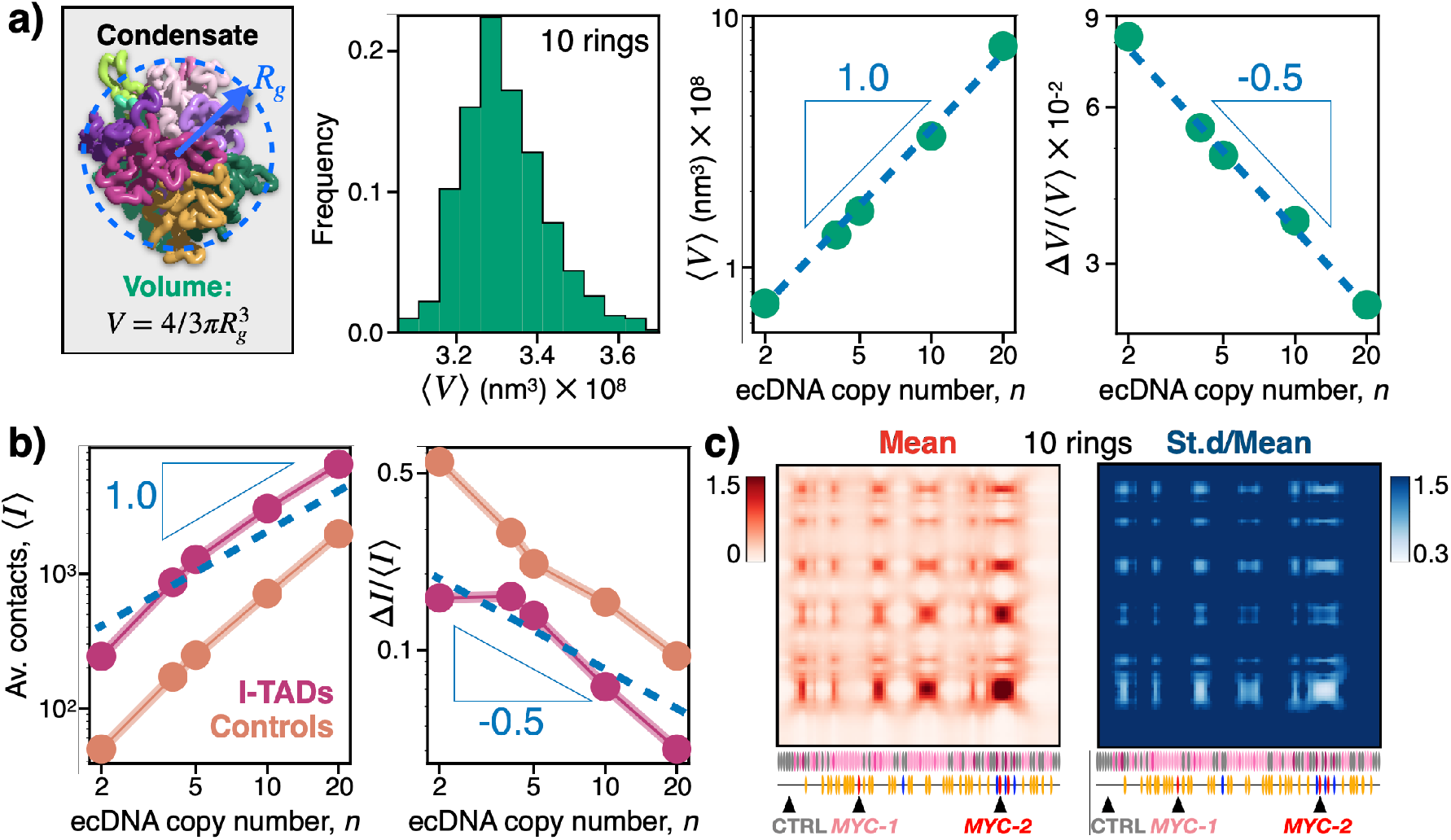
EcDNA condensates obey robust scaling laws with copy number. **a)** Condensate volume scales with ecDNA copy number, *n*. Left, representative 3D snapshot of a 10-ring condensate and definition of its effective volume, 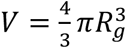, where *R_g_* is the condensate gyration radius. Centre-left, *V* has a broad distribution across single model conformations, shown here for *n*=10. Centre-right and right, the average condensate volume, ⟨*V*⟩, and its relative fluctuation, Δ*V*/⟨*V*⟩, scale as a power law with *n*, as expected from polymer physics. Their exponent is respectively around 1 and -1/2, showing that the relative fluctuations scale to zero as *n* grows. **b)** Left, the average number of contacts within I-TADs (purple) and size-matched control regions outside I-TADs (orange) also scales as a power law in *n*, approximately linear. Yet, I-TADs contacts are one order of magnitude higher that those outside I-TADs. Right, the relative fluctuation, Δ*I*/⟨*I*⟩, decreases approximately as *n*^−1/2^ in both cases, but remains markedly lower for I-TADs than for control regions. **c)** Maps of the average and relative fluctuations of total *in-trans* contacts, shown here for *n*=10, reveal that I-TADs corresponds to ecDNA genomic locations with markedly reduced relative contact fluctuations. Hence, the formation of I-TADs within large condensates stabilizes specific regulatory contacts, as reflected by their scaling behaviours with *n*.

Scaling laws also govern the organisation of contacts within ecDNA condensates, particularly within I-TADs. Under the hypothesis that contacts are overall independent across different rings, the Central Limit Theorem poses that their average number and relative fluctuations scale as a power law with *n*, with exponent equal to 1 and to -1/2 respectively. Indeed, we find that the average number of contacts in single I-TADs, ⟨*I*⟩, increases approximately linearly with copy number (**Fig. 3b**, left), reflecting two main underlying physical constraints: *(i)* copy-number enrichment, which linearly increases the number of potential interacting partners, and *(ii)* steric hindrance, which limits each ring to interacting with only a finite subset of neighbours. Size-matched control regions located outside I-TADs exhibit a similar scaling exponent, yet importantly their absolute contact levels remain nearly an order of magnitude lower, underscoring the selective, non-linear boost of contacts established within I-TADs by condensation (**Supplementary Fig. 4b**). Conversely, the relative fluctuations of I-TAD contacts, Δ*I*/⟨*I*⟩, decay approximately as *n*^−1/2^ (**Fig. 3b**, right), again consistent with Central-Limit averaging in larger assemblies. Control regions show a comparable exponent within our numerical accuracy but remain roughly an order of magnitude higher. Thus, increasing condensate size not only boosts I-TAD interactions but also suppresses their noise, producing mechanically and thermodynamically stabilised contact domains.

This behaviour is directly visible in maps of the relative fluctuations of total contacts (**Fig. 3c**), which exhibit pronounced minima in the standard-deviation-to-mean ratio precisely at I-TAD genomic positions. These dips reflect the local stiffening of the contact network imposed by BRD4-mediated bridging, establishing I-TADs as structurally stabilised modules within an otherwise heterogeneous polymer assembly.

For comparison, we also examined the dispersed phase, where no shared condensate is formed. In that regime, average *in-trans* contacts still increase with copy number and their relative fluctuations still decrease with *n*, as expected from simple counting statistics (**Supplementary Fig. 5**). However, contact levels remain about two orders of magnitude lower, and relative fluctuations about two orders of magnitude higher, than in the condensed phase. Thus, although copy-number scaling of contacts is also present in the dispersed phase, its effects remain quantitatively weak because contacts are sparse and noisy. Condensation instead shifts the system into a distinct high-contact, low-noise regime in which I-TADs emerge and become both enriched and stabilised.

To check the robustness of these scaling behaviours, we considered model variants where polymer-binder affinities are varied over one order of magnitude in the weak-biochemical energy range, binder-binder interactions are changed, or the sequence organisation of binding sites is modified. Yet, as expected from polymer physics, same scaling behaviours are found (**Methods**). More broadly, these predictions are in line with recent experiments showing that ecDNA oncogene expression and its standard deviation grow linearly with copy number^11,24^ and are boosted when ecDNAs form hubs^15,16^.

Taken together, these analyses show that ecDNA condensates, despite their broad structural heterogeneity, form stable, specific domains enriched for contacts, which follow simple and robust scaling laws with copy number. As condensates grow, contact strength within I-TADs increases linearly with ring number, while relative fluctuations decrease. By contrast, equivalently sized regions outside I-TADs remain markedly weaker and noisier, implying that the formation of hubs has a dual effect: boosting and stabilising contacts, particularly within I-TADs. Because I-TADs in COLO320-DM overlap enhancer-rich regulatory regions^15,16,18^ (**Fig. 3c**), these results indicate that high-copy number condensates create strong and stable regulatory environments for selected oncogenes, such as the *PVT1-MYC* fusions. This may provide a physical explanation for why individual tumour cells often harbour dozens to hundreds of ecDNAs: increasing ecDNA copy number enforces universal scaling behaviours that selectively boost oncogene regulatory contacts within I-TADs and suppress their relative fluctuations, potentially conferring a selective advantage to cancer cells.

### Structural variability of individual ecDNAs in a condensate

Next, we investigated how condensate-level heterogeneity arises from the variability in the conformations of their individual ecDNA rings, and to what extent distinct rings share or diverge in their 3D architecture. To this aim, we computed Pearson correlations, r, between pairs of single-ring distance matrices, separately *in-cis* and *in-trans*, in the model condensed phase. The resulting distributions (*cis-vs-cis, trans-vs-trans, cis-vs-trans*) are broad, with standard deviations ranging from about 30% to 90% of their average values, indicating that single ecDNAs sample a wide ensemble of thermodynamically allowed conformations (**Fig. 4a; Methods**).

**Figure 4.**
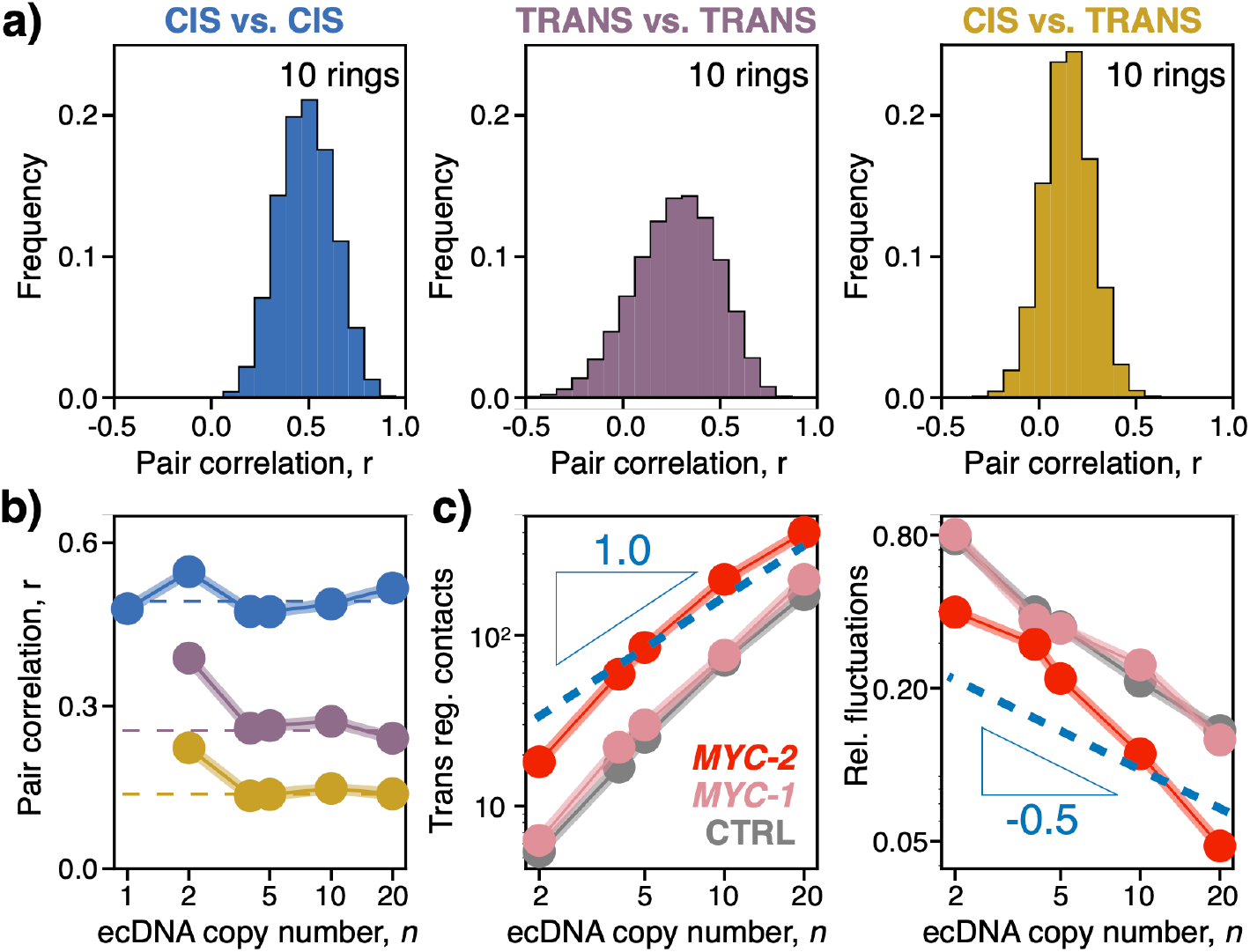
Variability of contact patterns of individual ecDNA polymers in a condensate. **a)** Distributions of Pearson correlation coefficients, r, between pairs of single-ring distance matrices in a 10-ring condensate, for cis-versus-cis (left), trans-versus-trans (centre), and cis-versus-trans (right) comparisons. All three distributions are broad, revealing substantial variability in the 3D conformations of individual ecDNA rings and in their pairwise spatial organisation. **b)** Average pairwise correlations as a function of ecDNA copy number, *n*. Cis-versus-cis correlations remain comparatively high (r=0.49), whereas trans-versus-trans and cis-versus-trans correlations plateau at lower yet still positive values (r=0.27 and r=0.15, respectively), indicating that thermal fluctuations generate broad single-molecule 3D heterogeneity, yet preserve a conserved interaction scaffold across condensates. **c)** Scaling of gene-specific *in-trans* regulatory contacts and their relative fluctuations. Left, total regulatory contacts involving the *PVT1-MYC* fusion (*MYC-2*, red), the canonical *MYC* copy (*MYC-1*, pink) and a control locus outside I-TADs (CTRL, grey) all increase approximately linearly with *n*, but remain about one order of magnitude higher for *MYC-2*, consistent with its localization within I-TADs. Right, the relative fluctuations of the same quantities decrease with *n*, with *MYC-2* displaying markedly lower variability than *MYC-1* and CTRL, which exhibit comparable behaviours. Thus, despite broad single-molecule heterogeneity, loci embedded within I-TADs acquire stronger and more stable regulatory interactions.

Despite this broad conformational degeneracy, all correlation distributions in the condensed phase are centred at non-zero average values. As the number of rings *n* increases, the average *cis-vs-cis* correlation remains comparatively high, r=0.49, whereas the *trans-vs-trans* correlation mildly decreases with *n* to plateau at r=0.27, as much as *cis-vs-trans* correlations plateauing at r=0.15 (**Fig. 4b**). Those values indicate that, as intuitively expected, contacts within a ring remain comparatively stable under thermal fluctuations, while intermolecular contacts fluctuate more strongly due to the larger accessible configurational space. Nevertheless, the persistence of non-zero average correlations highlight the existence of a common interaction scaffold, shaped by the genomic arrangement of BRD4 binding sites and by the emergence of I-TADs in the condensed state. Conversely, in the dispersed phase, correlations involving trans matrices are trivially lost, consistent with the absence of a shared interaction scaffold. Analogously, a null control obtained from randomised distance matrices is centred around zero (**Supplementary Fig. 6**). Thus, despite substantial thermal variability, ecDNA condensates retain a reproducible interaction pattern at the single-molecule level. This conserved scaffold provides the structural basis for specific and robust oncogene regulatory contacts within I-TADs and may help explain the preferential amplification of selected oncogenes in COLO320-DM cells^15,16^.

### **I-** TADs boost and stabilise *MYC* regulatory contacts

To determine how single-ring conformational variability affects the regulatory landscape of specific oncogenic loci, we next focused on the *PVT1-MYC* fusions (*MYC-2*) and the canonical *MYC* copy (*MYC-1*), which lie respectively inside and outside I-TADs. Consistent with the scaling behaviours described above, the total number of *in-trans* regulatory contacts involving *MYC-2* grows approximately linearly with the number of ecDNA rings *n*, as expected from copy-number effects (**Fig. 4c**, left). *MYC-1* and a control locus (CTRL) located outside I-TADs display a similar scaling exponent (**Fig. 4c** and **Fig. 5a; Methods**), but their absolute contact levels remain about one order of magnitude lower than those of *MYC-2*, indicating that loci embedded within I-TADs most efficiently recruit the regulatory interactions within the condensate.

**Figure 5.**
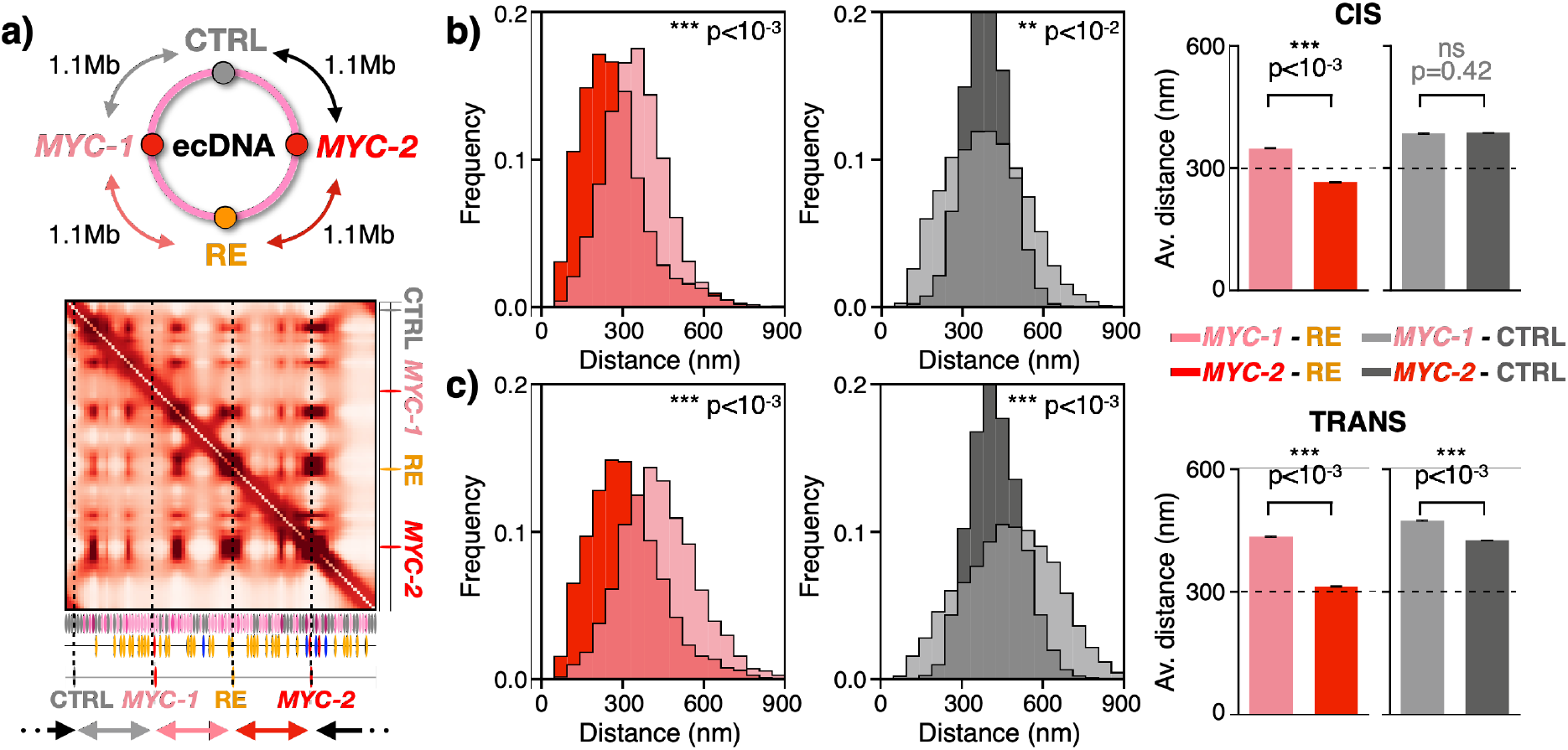
Variability of *MYC* interactions with its regulatory elements. **a)** Total contact map for a representative condensate with *n*=10 rings. The schematic above highlights the relative genomic positions of the canonical *MYC* copy (*MYC-1*), a *PVT1-MYC* fusion (*MYC-2*), a regulatory element (RE) located within an I-TAD, and a control locus (CTRL) outside I-TADs. RE and CTRL are both positioned at comparable genomic distance (≈1.1 Mb) from *MYC-1* and *MYC-2*. **b)** *In-cis* distance distributions and average-distance bar plots for *MYC-1*–RE (pink) and *MYC-2*–RE (red), compared with the corresponding distances to CTRL (gray shades). *MYC-2* is significantly closer to RE than *MYC-1*, whereas distances to CTRL are comparable for both loci. **c)** Same as **b)** for *in-trans* distances computed with the closest neighbouring ring. *MYC-2* remains significantly closer to RE than *MYC-1*, whereas the *MYC-1*– RE distance is comparable to the *MYC-1*–CTRL distance. Thus, although single-gene interactions remain highly dynamic, loci embedded within I-TADs, such as *MYC-2*, experience more stable and less variable regulatory environments than loci outside these domains, such as *MYC-1* and CTRL. P-values for distance distributions and bar plots were computed using, respectively, two-sided Mann-Whitney U-tests and two-sided Student’s t-tests.

Contact stability follows a complementary trend: the relative fluctuations of regulatory contacts involving *MYC-2*, *MYC-1*, and CTRL all decrease with *n* as a power law with an exponent around -0.5, consistent with fluctuations averaging out in larger polymer assemblies (**Fig. 4c**, right). However, *MYC-2* exhibits markedly lower relative fluctuations than *MYC-1* and CTRL, by about one order of magnitude, reflecting the stabilising effect of its incorporation into I-TADs. Thus, as condensates grow, *MYC-2* not only acquires substantially more regulatory contacts than *MYC-1*, but does so in a much less noisy environment. These results highlight that I-TAD membership, rather than genomic position alone, governs both the strength and the stability of enhancer-oncogene communication within ecDNA condensates.

We then asked whether this difference is also reflected in direct 3D spatial proximity to regulatory elements. To this end, we considered a regulatory element (RE) located at comparable genomic distance (≈1.1Mb) from *MYC-1* and *MYC-2*, and compared it with the control locus CTRL, positioned outside I-TADs and also genomically equally distant from them (**Fig. 5a**; **Methods**). Despite comparable genomic separations, RE is significantly closer in 3D space to *MYC-2* than to *MYC-1*, both *in-cis* and *in-trans* (**Fig. 5b,c**). In particular, the average *in-cis* distance between *MYC-2* and RE is 260 nm, approximately 30% shorter than the corresponding *MYC-1*–RE distance of 350 nm (**Fig. 5b**), reinforcing the conclusion that I-TAD membership, rather than linear genomic distance on the ecDNA sequence, determines regulatory proximity. Moreover, the average *in-trans* distance between *MYC-2* and RE on the closest neighbouring ring is 310 nm, only about 20% larger than their *in-cis* distance. By contrast, the average *in-trans* distance between *MYC-1* and RE is 440 nm, comparable to the *MYC-1*–CTRL distance *in-trans* and larger than the corresponding distance *in-cis* (**Fig. 5c**), consistent with the exclusion of *MYC-1* from I-TADs.

Although all distance distributions remain broad (**Fig. 5b,c**), as expected for thermally fluctuating polymers and consistent with experimental observations^16,49^, their shifted average values reveal a systematic reshaping of the regulatory contact landscape inside condensates. I-TAD membership places *MYC-2* within regions of elevated and stabilised local contact probability, enabling more frequent and persistent regulatory encounters than those accessible to *MYC-1* outside these domains. This structural bias provides a mechanistic explanation for the experimentally observed preferential regulatory engagement of *PVT1-MYC* fusions within ecDNA condensates.

## Discussion

Our study shows that ecDNA condensates in COLO320-DM cancer cells operate as self-organising polymer assemblies that selectively shape oncogene regulatory architectures. Using a minimal yet mechanistically grounded polymer model, we find that BRD4-like complexes drive a phase transition that compacts multiple ecDNA rings into a single condensate. Although thermal fluctuations generate substantial variability in the 3D conformations of individual rings and in their pairwise spatial distances, condensates retain a conserved structural scaffold determined by the ecDNA binding-site landscape and by the underlying thermodynamic mechanism of condensate assembly.

Within this scaffold, condensates establish enriched contact domains *in-cis* (TADs) and a combinatorial number of contact domains *in-trans* (I-TADs), whose genomic positions are encoded by the genetic arrangement of BRD4 binding sites along the ecDNA sequence. At the same time, key condensate properties follow robust polymer-scaling behaviours with ecDNA copy number, including condensate volume, regulatory contact strength and contact fluctuations. In particular, loci embedded within I-TADs experience both comparatively amplified and markedly stabilised regulatory contacts as ecDNA copy number in the condensate increases. These behaviours are universal as they do not depend on the minute details of the interaction potential or specificity of the involved molecular factors. These physical mechanisms provide, thus, a robust route by which high copy-number ecDNA condensates generate strong and stable enhancer-oncogene communication, despite their inherently high structural heterogeneity (**Fig. 6**).

**Figure 6.**
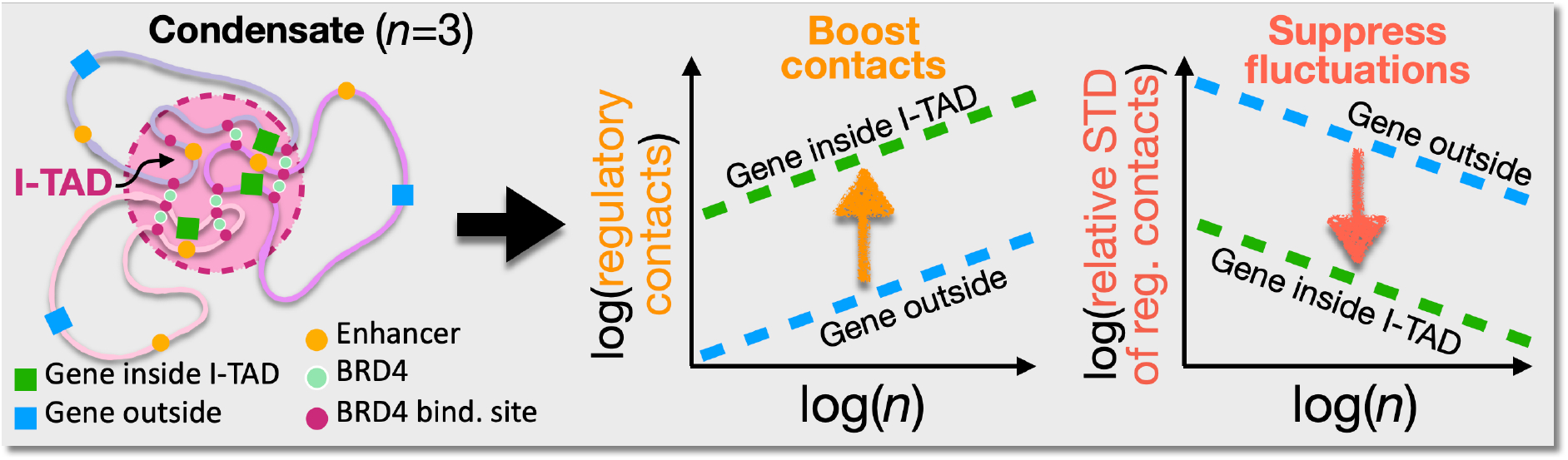
Graphical summary. The assembly of high copy-number ecDNA condensates, despite broad single-molecule structural heterogeneity, enforces universal scaling laws whereby oncogene regulatory contacts within I-TADs are boosted and their relative fluctuations suppressed, stabilising specific oncogene activation and potentially conferring a selective advantage to the cell.

Our analysis also clarifies why only specific oncogenes, rather than all ecDNA-encoded genes, gain regulatory enhancement. In COLO320-DM cells, the preferential activation of *PVT1-MYC* fusions arises not from their genomic distance to regulatory elements, but from their systematic localisation within I-TADs. By contrast, the canonical *MYC* copy, positioned outside I-TADs, is disproportionally less amplified. This distinction points to a more general principle: within ecDNA condensates, the 3D spatial organisation of loci, rather than linear genomic sequence alone, is a key determinant of regulatory output.

The model we consider is intentionally minimal and lacks several ingredients, such as chromosomal context, multiple ecDNA species, nuclear landmarks, and active folding processes, which are likely to play relevant roles in real cells. Nevertheless, the mechanisms we uncover are robust, as they are rooted in general thermodynamic and polymer physics principles, and could have broader implications for understanding ecDNA function in cancer evolution. The scaling-based enhancement of regulatory contacts and suppression of noise suggests that tumour cells harbouring ecDNA condensates achieve more persistent oncogene transcription, consistent with experimental observations that COLO320-DM hubs contain dozens of highly transcribing ecDNAs^10,15,16,28^, potentially contributing to the selective advantage of high-copy-number ecDNA states.

Taken together, our results provide quantitative predictions on how ecDNA copy number variation shapes 3D structural heterogeneity, regulatory contacts and oncogene activation, which can be tested, e.g., by emerging single-molecule ecDNA tracing approaches^50^. The universal scaling behaviours identified here offer a predictive biophysical basis for interpreting ecDNA-driven phenotypes across tumour contexts, including intratumoral heterogeneity, tumour evolution and therapeutic response. More generally, they may help connect the physical organisation of ecDNAs to their broader functional and clinical consequences.

## DATA AVAILABLITY

Raw molecular dynamics simulation data generated in this study will be available on Zenodo.

## CODE AVAILABLITY

All codes used in this work are based on standard, publicly available software packages, as detailed in the Methods. Molecular dynamics simulations were performed using LAMMPS (12 Dec 2018 release). 3D renderings were generated using POV-Ray (version 3.7). Simulation codes, input files and analysis scripts will be available on Zenodo.

## ACKNOWLEDGMENTS

MN acknowledges support from MUR PRIN 2022 2022R8YXMR CUP E53D23001810006, MUR PRIN 2022 PNRR P2022JAYMH CUP E53D23018360001, and computer resources from INFN, CINECA, ENEA CRESCO/ ENEAGRID^51^ and Ibisco at the University of Naples.

## AUTHOR CONTRIBUTIONS

MN and MC designed the project. MN and MC developed the modelling. SK and MC ran computer simulations and performed data analyses with help from SG. MN and MC wrote the manuscript with inputs from SK and SG.

## METHODS

### Polymer model of COLO320-DM ecDNAs

We studied a 4.328 Mb *MYC*-harbouring ecDNA in COLO320-DM colorectal cancer cells, containing fragments derived mainly from chromosome 8 along with smaller segments from chromosomes 6, 13 and 16, and including a canonical *MYC* copy, *PVT1-MYC* fusions and mapped cis-regulatory elements^15,16^. To investigate its three-dimensional organisation, we used the Strings and Binders (SBS) model^30–34^, a coarse-grained polymer model of chromatin in which each ecDNA is represented as a self-avoiding polymer ring made of beads carrying binding sites for cognate diffusing molecular binders. In the present implementation, beads are of two types: inert beads, which interact only through polymer connectivity and steric exclusion, and BRD4-binding beads, which can be bridged by diffusing BRD4-like complexes and thereby generate both intra-ring and inter-ring interactions, referred to hereafter as *in-cis* and *in-trans* contacts, respectively.

To build the polymer sequence, the experimentally reported ecDNA genomic fragments^15^ were aligned to the reference genome and coarse-grained at 50 kb resolution, assigning one polymer bead to each retained genomic bin (**Supplementary Fig. 1**). Beads with only minimal overlap with an ecDNA fragment (<8kb) were excluded unless they corresponded to biologically relevant loci such as *MYC* or *PVT1*. The order of beads along the polymer ring reproduced the experimentally inferred fragment arrangement of the COLO320-DM ecDNA^15^. BRD4-binding sites were mapped from BRD4 ChIP-seq data^15^ binned at the same resolution. When a ChIP-seq peak overlapped genomic coordinates present in more than one ecDNA fragment, the corresponding signal was divided by the number of overlapping fragments before assignment to the model, to avoid artificial overcounting. The BRD4 signal amplitude was then discretised into five equally spaced affinity classes, which in the model span the weak biochemical range 2.25-6.25 k_B_T. The different copies of *MYC* and *PVT1*, together with the cis-regulatory elements, were mapped onto the same polymer chain and subsequently used as genomic viewpoints in the analyses. A linear scheme of the model is in **Fig. 1a**.

In the main case study, the diffusing binders represent BRD4-containing complexes rather than isolated BRD4 molecules. To account in a minimal way for stabilising cofactor-mediated interactions, binders were assigned a weak mutual affinity. In the model, this binder-binder interaction was set to 3.1 k_B_T, in line with in vitro measurements of IDR-mediated droplet formation^35^ and previous polymer studies^33,36–38^. Model variants spanning affinities up to 10 k_B_T yielded similar contact patterns, confirming the robustness of the main results. This model focuses specifically on ecDNA self-organization and neglects other chromosomes and nuclear landmarks, so as to isolate the minimal physical ingredients sufficient to account for ecDNA condensation and the emergence of specific *in-trans* associated domains.

### Molecular Dynamics simulations

We performed Molecular Dynamics (MD) simulations with Langevin dynamics in a cubic box with periodic boundary conditions, using standard coarse-grained polymer interaction potentials^36,52,53^. Consecutive beads along each ring were connected by a finitely extensible nonlinear elastic (FENE) potential, excluded-volume interactions were modeled by the Weeks-Chandler-Andersen (WCA) repulsion, and specific attractions between BRD4-binding beads and cognate binders were implemented using a standard truncated and shifted Lennard-Jones potential. Simulations were performed in LAMMPS^54^ starting from random initial conditions consisting of self-avoiding ring conformations and randomly distributed binders. Systems were equilibrated for up to 10^8^ MD timesteps and configurations were sampled at steady state. All simulations parameters were set as in^18^.

We explored systems with ecDNA copy number, *n*, ranging from 1 to 20. The case *n* = 10 was used as a representative reference throughout, consistent with experimental observations that COLO320-DM ecDNA hubs typically contain on the order of ten rings^15^. The simulation box size was fixed to 40σ, where σ is the bead diameter, which was sufficiently large to avoid detectable finite-size effects across all investigated conditions. In each simulation, the number of rings was fixed and only the binder concentration was varied to probe the phase behavior of the system.

Physical units were assigned by calibrating σ against microscopy measurements of COLO320-DM ecDNA hubs^15^. Matching the experimentally reported median size of ecDNA clusters (≈700 nm) with the median steady-state diameter of a condensate in the model (≈11σ, estimated as twice the gyration radius) yielded σ ≈ 64 nm^18^. This value should be interpreted as an effective coarse-graining scale, which may vary with local chromatin context (e.g., compaction level, epigenetic state, nucleosome organization, or cell type), thereby contributing to differences in bead-size estimates across studies. Binder concentrations were converted into molar units as [BRD4]=P/(V_B_N_A_), where P is the number of binders, V_B_ the simulation box volume, and N_A_ Avogadro’s number. In this study we explored concentrations spanning four orders of magnitude, from 10^0^ to 10^3^ nM.

As [BRD4] increases, the system undergoes a transition from a dispersed state, in which ecDNA rings are randomly distributed in the available volume, to a condensed state, in which they phase-separate into a single cluster. In the model, this transition is marked, e.g., by a sharp decrease in the gyration radius of the whole system, *R*_g_ (**Fig. 1c**), or in the distance between rings’ mass centres (**Supplementary Fig. 2**). Unless otherwise stated, analyses were performed in the condensed regime, using above-threshold binder concentrations. In the cases considered here, we used [BRD4] = 50, 50, 150, 300, 500, 1000 nM for systems with *n* = 1, 2, 4, 5, 10, 20, respectively. Dispersed-phase measurements were obtained by evaluating the same observables at sub-threshold [BRD4]. Importantly, as dictated by physics, the model emergent conformations are determined primarily by the underlying thermodynamic state of the system, rather than by the precise value of [BRD4] within a given regime. For each parameter set, we collected up to 10^3^ independent equilibrated conformations for downstream analyses.

### 3D size of model ecDNA condensates

For each equilibrated conformation in the condensate regime, we computed the gyration radius, *R*_g_, of the whole ecDNA aggregate by considering all polymer beads from all rings. We then defined an effective condensate volume as 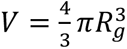. For each copy number *n*, we measured the distribution of *V* across independent conformations, together with its mean, ⟨*V*⟩, and relative fluctuation, Δ*V*/⟨*V*⟩, where Δ*V* = (⟨*V*^2^⟩ − ⟨*V*⟩^2^)^1/2^. Their scaling with *n* is reported in **Fig. 3a**.

### Contact and distance matrices of model ecDNAs

Two polymer beads were defined to be in contact if their Euclidean distance was below a spatial threshold, *t_c_*, consistent with standard practice in coarse-grained chromatin polymer models^31,36^. We set *t_c_* = 200 nm, in line with contact thresholds used in microscopy-based chromatin proximity analyses^55,56^, yet we also checked that results are robust to variations within the 100-300 nm range. To calculate ensemble-averaged contact matrices (**Fig. 2b**), we reported contact observables normalized by the number of rings *n*. Let *C*(*i_k_*, *j*_ℎ_) denote the contact frequency between locus *i* on ring *k* and locus *j* on ring *h*. The ensemble-averaged *in-cis*, *in-trans*, and total contact matrices were defined as:

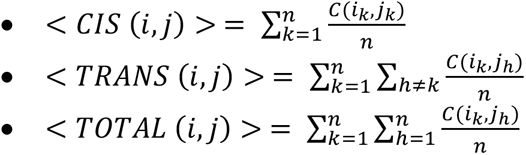

Distance matrices were computed analogously by replacing contacts with spatial distances. To avoid the trivial reduction of mean *in-trans* distances caused by increased packing at larger copy number, in **Fig. 2b** each ring in each conformation was paired with its nearest distinct neighbour in 3D space, as identified from the centre-of-mass distance between rings, and the corresponding pairwise inter-ring distance matrix was computed for that pair only.

Variability of total contacts *in-trans* as reported in **Fig. 3c** was quantified as follows: in each sampled conformation, and for each ring in the condensate, we computed its total *in-trans* contact matrix by summing the contacts between that ring and all other *n*-1 rings. Pooling all rings across ≈10^3^ independent conformations yielded an ensemble of total *in-trans* contact matrices. The mean matrix was defined as the element-wise average over this ensemble, and the corresponding variability map as the element-wise ratio of standard deviation to mean. The same total *in-trans* ring contact matrices were consistently used for the *in-trans* contact analyses reported in **Figs. 3b,4c**.

### I-TAD calling and control regions

I-TADs were identified from the ensemble-averaged total *in-trans* contact matrix using a trans-insulation-like quantity that we termed the I-TAD enrichment score (**Supplementary Fig. 4a**). For each putative boundary at genomic position *i* + 1/2, we defined a left block *L_i_* and a right block *R_i_*, each spanning *w* bins, and computed:

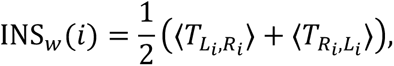

where *T* is the total *in-trans* contact matrix and ⟨*T_Li_*_,*R*_*_i_*⟩ denotes the mean matrix entry between the two blocks. To enhance local trans-contact enrichment against a broader surrounding background, we computed two tracks: a narrow one with *w*_narrow_ = 2 bins and a wide one with *w*_wide_ = 4 bins. We also checked that results were robust to varying these parameters over a broader range up to *w*_narrow_ = 5 and *w*_wide_ = 10 bins. We then defined the I-TAD enrichment score as:

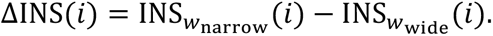

Positive local maxima of ΔINS(*i*) were taken as candidate I-TAD centres. For each selected peak, the I-TAD width was defined as the full width at half prominence of the corresponding ΔINS peak, i.e. the width measured at half of the peak height above its local background set by the neighbouring flanking minima. The resulting interval, rounded to the nearest bin, defined the genomic extent of the I-TAD and the corresponding square block in the total *in-trans* contact matrix. Size-matched control regions were obtained by rigidly shifting each called I-TAD block by -7 bins along both matrix axes, towards lower genomic coordinates, obtaining windows that did not overlap the called I-TADs and that were then used as the null comparison for I-TAD contact strength and variability (**Supplementary Fig. 4a**).

### Scaling analyses

For each copy number *n*, we quantified both the mean and the fluctuations of condensate- and domain-level observables across independent equilibrated conformations. For condensate size, the observables were ⟨*V*⟩ and Δ*V*/⟨*V*⟩, as defined above (**Fig. 3a**). For contact domains, we computed, for each I-TAD or matched control region, the total *in-trans* contact count, *I*, by summing the entries of the per-ring total *in-trans* contact matrix within the corresponding I-TAD or control square block. We then evaluated the ensemble mean, ⟨*I*⟩, and the relative fluctuation, Δ*I*/⟨*I*⟩, where Δ*I* = (⟨*I*^2^⟩ − ⟨*I*⟩^2^)^1/2^(**Fig. 3b, Supplementary Fig. 4b**). These observables were also measured below the phase-separation threshold to compare the scaling behaviour in the dispersed and condensed regimes (**Supplementary Fig. 5**). The same definitions were used for the gene-based trans-contact observables in **Fig. 4c**, where the sum was taken over matrix entries connecting a selected genomic viewpoint (*MYC-1*, *MYC-2*, and CTRL; see below) to bins annotated as regulatory elements on the other rings.

Scaling exponents were obtained by fitting the dependence on copy number with power laws of the form *y*(*n*) = *an^b^*. Fits were performed in Python (v.3.8) using standard SciPy routines. In particular, we tested the expectation that extensive quantities, such as condensate volume and total *in-trans* contacts, scale approximately linearly with *n*, whereas their fluctuations scale as *n*^-1/2^, as discussed in the text. Unless otherwise specified, fits were performed in log-log coordinates over the full range of simulated *n* values shown in the corresponding figure. Statistical errors on the mean data points shown in the figures are contained within the symbol size.

### Pairwise structural correlations

To quantify single-molecule structural heterogeneity, we computed Pearson correlations, r, between pairs of single-molecule distance matrices in the model condensed state. We considered three classes of comparisons: cis-cis, trans-trans and cis-trans. For each class, we analysed the full distribution of r values together with its mean and standard deviation, as reported in **Fig. 4a,b**. The same analysis was repeated for *n*=1,2,4,5,10,20. For correlations involving *in-cis* matrices, the main diagonal was masked before flattening to avoid trivial inflation from self-distances. As a null control, the entries of single-molecule distance matrices were randomly reshuffled, thereby preserving the distribution of distances while destroying their spatial organization. Correlation distributions obtained from these reshuffled matrices were centred around zero, as expected for matrices lacking reproducible structure (**Supplementary Fig. 6**). We further verified that replacing Pearson with Spearman rank correlation led to similar conclusions.

### Gene-specific contact and distance analyses

We analysed a canonical *MYC* copy (*MYC-1*), a *PVT1-MYC* fusion (*MYC-2*), a regulatory element (RE) and a control locus (CTRL). RE and CTRL were chosen to lie at comparable genomic distance (≈1.1Mb) from *MYC-1* and *MYC-2*, with RE located inside an I-TAD and CTRL outside (**Fig. 5a**).

For the distance analyses in **Fig. 5b,c**, *in-cis* distances were measured between loci located on the same ring. *In-trans* distances were defined using the closest distinct neighbouring ring in 3D space, identified from the centre-of-mass distance between rings. For a given pair of loci, we quantified the shortest inter-ring 3D distance, in analogy with ecDNA microscopy-based procedures in which each locus is assigned the minimal distance to the nearest relevant target^16,49^. The resulting distance distributions were used to compare the spatial proximity of *MYC-1* and *MYC-2* to regulatory versus control loci *in-cis* and *in-trans*.

For the barplot summaries shown in **Fig. 5b,c**, mean locus-locus distances were obtained by averaging the corresponding cis or minimal-trans distances across sampled conformations. Statistical significance of the full distance distributions was assessed using a two-sided Mann-Whitney U-test, whereas differences between mean values were assessed using a two-sided Student’s t-test, as indicated in the corresponding figure captions.

### Robustness analyses

To assess robustness, we examined model variants previously explored for the same ecDNA system^18^. These included alternative ranges of DNA-binder affinity (4.25-8.25 and 6.25-10.25 k_B_T), binder-binder affinities ranging from 0 to 10 k_B_T, a finer 10 kb coarse-grained model representation, and a variant in which BRD4 affinities were assigned from raw ChIP-seq signal without correction for overlapping ecDNA fragments. Across all these variants, the main *in-cis*, *in-trans*, and total contact patterns were preserved, as well as the genomic positions of the identified I-TADs, indicating that the main structural conclusions of the model are robust to specific parameter choices.

